# The molecular basis of the synergistic toxicity of Ni and Cu, common environmental co-contaminants

**DOI:** 10.1101/2025.08.18.670860

**Authors:** Linda Darwiche, Carlos A. Rodriguez-Bornot, Rebecca A. Ingrassia, Max J. Loccisano, Gray Waldschmidt, Jennifer L. Goff

## Abstract

Heavy metals are ubiquitous in the environment due to both natural processes and anthropogenic activities. Nickel (Ni) and copper (Cu) commonly co-occur in contaminated environments, yet most toxicity studies focus on individual metals. We investigated the combined toxicity of Ni and Cu in *Escherichia coli* using environmentally relevant concentrations of each. While each metal alone caused minimal growth inhibition, their combination was synergistically toxic. Transcriptomic and metabolomic analyses revealed unique alterations in gene expression and metabolites during the combined metal treatment. Key pathways uniquely impacted by the combined metal exposure included sulfur assimilation, cysteine biosynthesis, and the tricarboxylic acid cycle. Many of these responses appeared to be linked to dysregulation of iron-sulfur (Fe-S) cluster metabolism. We observed increased expression of the genes encoding ISC Fe-S cluster assembly machinery only during metal co-treatment. Growth experiments with deletion mutants confirmed that the ISC machinery was required for survival only under the combined metal stress. We also observed the activation of a sulfur starvation response during the combined metal stress that was consistent with increased sulfur demand for Fe-S cluster biosynthesis. Deletion of *cysK*, encoding cysteine synthase, impaired growth only under combined metal exposure. Because Fe-S clusters are universal across microbial taxa, the common co-occurrence of Ni and Cu in the environment represents a widespread and underrecognized threat to microbial life and the ecosystem processes they sustain. Our findings highlight the need to further assess the effects of metal mixtures, which can trigger emergent stress responses not yet predictable from single-metal exposures.

**IMPORTANCE:** Many environments are contaminated by metals. These metals are toxic to the microorganisms that inhabit these environments and carry out important ecosystem services. While much is known about bacterial responses to single metal stress, in most contaminated environments, metals typically exist as mixtures. Nickel (Ni) and copper (Cu) are common co-contaminants. We tested Ni and Cu in combination to shed light on the mechanism behind their synergistic toxicity in the model bacterium *Escherichia coli* K-12 using a novel, multi-omics approach. We found that the two metals in combination are likely disrupting iron-sulfur (Fe-S) clusters. Since Fe-S clusters are ubiquitous across microbial taxa and critical for microbial metabolism, this suggests that these two common co-contaminants may be toxic to diverse microorganisms.

## INTRODUCTION

Globally, heavy metal contamination has increased significantly since the Industrial Revolution began in the 19th century (1). Although heavy metals are needed in trace amounts for biological processes such as electron transport, metabolism, enzyme regulation (2), at higher concentrations, all heavy metals are toxic (3). High levels of heavy metals can be harmful to living organisms, including bacteria performing key ecosystem services (4–10).

The presence of heavy metals in the environment impacts bacterial biochemistry. Mechanisms of metal ion toxicity in bacteria include metalloenzyme mismetallation (*i.e.,* insertion of the incorrect metal cofactor) (11), antagonism of essential metal ion uptake (12), generation of reactive oxygen species (ROS) (13), and induction of disulfide stress (14). Bacteria have numerous resistance mechanisms that allow them to survive this heavy metal stress. These mechanisms are intrinsic or acquired by chromosomal mutations or through transferable genetic materials (15–19) and include increased expression of efflux pumps, decreased expression of uptake systems, increased expression of metal-binding proteins, and increased expression of metal-reducing oxidoreductases—among others (15). This body of literature describing mechanisms of heavy metal toxicity and resistance in bacteria is largely derived from studies conducted with individual metal exposures. However, in most environments, heavy metal contamination typically does not involve just one individual metal. Rather, heavy metals tend to exist as contaminant mixtures (20–22).

Several studies have highlighted the synergistic and antagonistic effects of metal combinations on bacterial growth (23–29) but did not examine the underlying molecular mechanisms of these effects. In a recent study, we utilized an environmental *Bacillus cereus* isolate—originating from a heavy metal-contaminated site—to examine the effects of a site-informed eight-metal mixture on cell physiology through a combination of proteomic and metabolomic analyses (30). We observed that the metal mixture elicited a unique physiological response that could not be predicted by the single metal exposures alone. In analyzing the proteome shifts across all stress conditions (metal mixture exposure and eight individual metal exposures), we found that 65% of differentially abundant proteins were differentially abundant only under the metal mixture stress. These findings highlighted the limitations of single metal stress experiments in fully understanding the mechanisms by which bacteria respond to heavy metal stress in their environment. However, disentangling metal interactions in a non-model bacterial system is challenging—further compounded by the difficulty in studying an eight-metal mixture. Thus, there is a need to develop a foundational, predictive understanding of metal interactions using simplified combinations of heavy metals in model organisms.

Nickel (Ni) and copper (Cu) are two commonly co-occurring heavy metals in the environment (31,32), often introduced through anthropogenic activities (33). While each metal, individually, has distinct toxicological effects, their interactions within bacterial systems remain underexplored. An early study described the synergistic toxicity of Ni and Cu in *Bacillus subtilis, Enterobacter aerogenes,* and *Nocardia corallina* (31). However, no mechanistic details were provided, and there was no follow-up on this work. Additionally, our recent study with the *B. cereus* isolate suggested that the synergism between Ni and Cu was a significant contributor to the overall toxicity of the eight-metal mixture (30). In this current study, we utilized environmentally relevant concentrations of Ni and Cu to confirm their synergistic toxicity in *Escherichia coli* K-12, our model system for this analysis. Utilizing a combination of transcriptomics, metabolomics, and screens of mutant strains, we examined how exposure to a combination of Ni and Cu impacts *E. coli* growth and functioning, providing a deeper understanding of their combined impact on microbial physiology.

## MATERIALS AND METHODS

### Bacterial strains

*Escherichia coli* K-12 BW25113 was used throughout the study, unless otherwise indicated. Strain BW25113 is the parental strain for the *E. coli* Keio Knockout Collection (34). Both the parental and mutant strains were purchased from Horizon (Cambridge, UK).

### Media Preparation

To prepare LB liquid medium, 10 g of tryptone (IBI Scientific, USA), 10 g of NaCl (VWR, USA), and 5 g of yeast extract (MP Biomedicals, USA) were dissolved in 950 mL of distilled water (H_2_O). The solution was mixed until the solutes are fully dissolved, then the final volume was adjusted to 1 L by adding more distilled water. The medium was sterilized by autoclaving. For preparing LB agar plates, 15 g of agar was added to the solution before autoclaving to allow the medium to solidify. MES Buffered Minimal Medium (MBMM) was modified from Rathnayake et al. (2013)(35). The full recipe can be found under **SUPPPORTING METHODS** section in the **SUPPORTING INFORMATION** document.

### Heavy Metals Stock Solutions

Nickel (II) (Ni) chloride hexahydrate (MP Biomedicals, USA) was used to prepare 50 mM stock solution using deionized water (DI). Copper (II) (Cu) chloride dihydrate (Fisher Scientific, USA) was used to prepare 50 mM stock solution using DI H_2_O.

### Bacteria and Growth Conditions

For experimental work, glycerol stocks of *Escherichia coli* K-12 BW25113 (the parent strain of the Keio collection) and Keio collection mutants (34) were streaked onto LB plates and grown overnight at 37°C. A full list of the strains used can be found in **Supplementary Table S1.** Two to three colonies were selected and transferred to a liquid medium and then incubated in a shaker incubator at 37°C for 24 hours under constant shaking to allow bacterial growth before transfer into experimental growth medium.

### Bacterial Growth Curves

To evaluate the effects of specific supplements on *Escherichia coli* growth under metal stress, wild-type *E. coli* was cultured overnight in LB medium at 37 °C with shaking at 200 rpm. The next day, cultures were diluted 1:200 into MBMM and subjected to treatment with 30 µM NiCl₂, 15 µM CuCl₂, or a combination of both metals. Supplement additions, if utilized, were performed immediately after metal treatment. The experimental design included a control (no metal), one of three metal treatments (30 µM Ni, 15 µM Cu, or 30 µM Ni and 15 µM Cu), and each metal treatment in combination with the supplement additions (if used).

A range of concentrations was tested for each supplement to identify the most effective dose in restoring growth under metal stress conditions. These concentrations were selected based on their concentrations in common bacterial growth media, such as LB. For cystine, ten concentrations (0.2, 0.4, 0.6, 0.8, 1.0, 1.2, 1.4, 1.6, 1.8 mM) were tested and 0.8 mM was selected as the working concentration. For sulfate, five concentrations (2, 5, 10, 15, 20 mM) were tested using magnesium sulfate heptahydrate and 20 mM was selected as the working concentration. For glutathione, four concentrations (1, 2, 3, 5 mM) were tested and 1 mM was selected as the working concentration. For methionine, five concentrations (3, 5, 7, 10, 12 mM) were tested and 7 mM was chosen as the working concentration. For histidine, ten concentrations (0.3, 0.4, 0.5, 0.6, 0.8, 1.0, 1.2, 1.4, 1.6, 1.8 mM) were evaluated and 0.3 mM was selected as the working concentration. Iron supplementation was delivered using iron (II) sulfate heptahydrate, chelated with citrate in a 3:2 ratio (Fe:citrate). Five concentrations of Fe²⁺ (3, 6, 12, 25, 50 mM) were tested.

Following treatment setup, 200 µL of each culture was transferred into wells of a 96-well microtiter plate. The microplate was placed in a BioTek Synergy HTX plate reader, which monitored the optical density (OD) at 600 nm (OD600) while shaking and incubating the cultures at 37°C. Readings were taken by the plate reader every 30 minutes for 24 hours.

### Sample Preparation for Transcriptomics and Metabolomics

For transcriptomic and metabolomic analyses, bacterial cultures were prepared as described above, with a 1:200 dilution in MBMM and grown until they reached an OD of 0.25-0.3 (mid-exponential phase). Cultures were treated for 2 hours with the following conditions: [1] Control (no treatment), [2] 30 µM Ni, [3] 15 µM Cu, [4] 30 µM Ni and 15 µM Cu. After treatment, cultures were centrifuged at 10,000 rpm for 5 minutes, the supernatant was discarded, and the cell pellets were washed twice with phosphate-buffered saline (PBS). The washed pellets were resuspended in 1 mL of PBS, transferred to 2 mL sterile Eppendorf tubes, and centrifuged again. For metabolomic analyses, the final pellet was frozen at -80°C until shipment for analysis. For transcriptomics, RNA was immediately extracted using the Zymo Quick-RNA Miniprep Kit, following the manufacturer’s protocols. RNA samples were immediately frozen at -80°C prior to further analysis.

### Library Preparation and RNA Sequencing

Library preparation and RNA sequencing were performed at Biomarker Technologies (BMKGENE) USA Inc. (Durham, NC, USA) as described in the **SUPPPORTING METHODS** section in the **SUPPORTING INFORMATION** document.

### Transcriptome Assembly and Analysis

The RNA-seq analysis was performed using the US Department of Energy’s KnowledgeBase (KBase)(36) bioinformatics platform. A standardized workflow incorporating HISAT2 (37), StringTie (38), and DESeq2 (39) was followed. First, the *Escherichia coli* K-12 reference genome (RefSeq: NC_000913) was imported for alignment. Next, raw sequencing reads were obtained from an online repository and organized into a structured RNA-seq sample set. Read quality was assessed using FastQC (40) to identify potential sequencing artifacts. The reads were then aligned to the reference genome using HISAT2. Following alignment, transcripts were assembled using StringTie to reconstruct gene structures. Differential expression analysis was performed using DESeq2 to identify genes with significant expression changes. Finally, a filtered differential expression matrix was generated to refine the dataset for downstream functional analysis.

### LC-MS Metabolomics

The liquid chromatography-mass spectrometry (LC-MS)-based untargeted metabolomic analysis was performed at Creative Proteomics (Shirley, NY, USA). Extraction and analysis methods are described in the **SUPPPORTING METHODS** section in the **SUPPORTING INFORMATION** document.

### Metabolomic Data Analysis

For the metabolomics analysis, p-values were initially calculated using a Welch’s t-test to compare metabolite concentration averages between each of the metal treatments and the control. This was followed by adjustment for multiple comparisons using the Benjamini-Hochberg procedure.

### ROS Assay

2’,7’-Dichlorodihydrofluorescein diacetate (H2DCFDA) is a non-fluorescent molecule. Once imported into the cell, it is deacetylated to dichlorofluorescin (DCFH). When DCFH is exposed to reactive oxygen species (ROS), the molecule is oxidized to the fluorescent dichlorofluorescein (DCF) (41). To prepare the probe, 5 mg of H2DCFDA was dissolved in 5 mL DMSO to prepare a 2 mM solution, which was stored at -20°C. To measure ROS production, an overnight culture of the wild-type *E. coli* K-12 was diluted 200-fold into MBMM and left to regrow at 37° in a shaking incubator until mid-log growth phase (OD600 of 0.25-0.3). Mid-log cultures were centrifuged at 5900 x g for 5 minutes. The pellet was resuspended in sterile MBMM. The H2DCFDA probe was added at a final concentration of 5 µM. The cells were incubated with the probe for 1 hour at room temperature in the absence of light to allow uptake of the probe into bacterial cells. After incubation, the culture was centrifuged and resuspended in sterile MBMM. The cells with the probe were then exposed to their respective treatments (30 μM Ni, 15 μM Cu, and the mixture of the two) and monitored for fluorescence signal in our incubating plate reader. The fluorescence signal was monitored for 12 hours with excitation and emission wavelengths of 485/20 and 528/20 nm, respectively, using the bottom light source. Measurements were recorded every 15 minutes.

### Spot Dilution Assay

Spot-plating was performed to evaluate metal sensitivity of the wild-type *E. coli* and selected mutants from the Keio collection **(Suppl. Table S1).** Overnight cultures were grown in LB medium at 37 °C with shaking at 200 rpm. The following day, cultures were diluted in sterile PBS to an OD600 of 0.45. Tenfold serial dilutions (10⁰ to 10⁻⁵) were prepared in PBS, and 5 µL of each dilution was spotted in duplicate onto MBMM agar supplemented with the following: (1) control (MES without metal supplementation), (2) 30 µM NiCl₂, (3) 15 µM CuCl₂, and (4) combined treatment with 30 µM NiCl₂ and 15 µM CuCl₂. Plates for the control and single-metal treatments were incubated at 37 °C for 24 hours, while plates containing the combined metal treatment were incubated at 37 °C for 6 days to allow detection of delayed growth phenotypes. Plates were photographed using a smartphone camera under consistent lighting conditions.

## RESULTS AND DISCUSSION

### Ni and Cu commonly co-occur in freshwater environments

We investigated the individual and combined effects of Ni and Cu on *E. coli* as these two metals are commonly detected together in polluted ecosystems, particularly in areas influenced by industrial discharge, mining activities, and agricultural runoff (42,43). We first re-analyzed a dataset of heavy metal concentrations in 239 global rivers and lakes from 1972 to 2017 (44). We performed a log-log linear regression to evaluate the power-law relationship between [Ni] and [Cu] in this dataset. Our analysis revealed a statistically significant, positive correlation between the concentrations of Cu and Ni **(Fig. 1A).**

**Figure 1.**
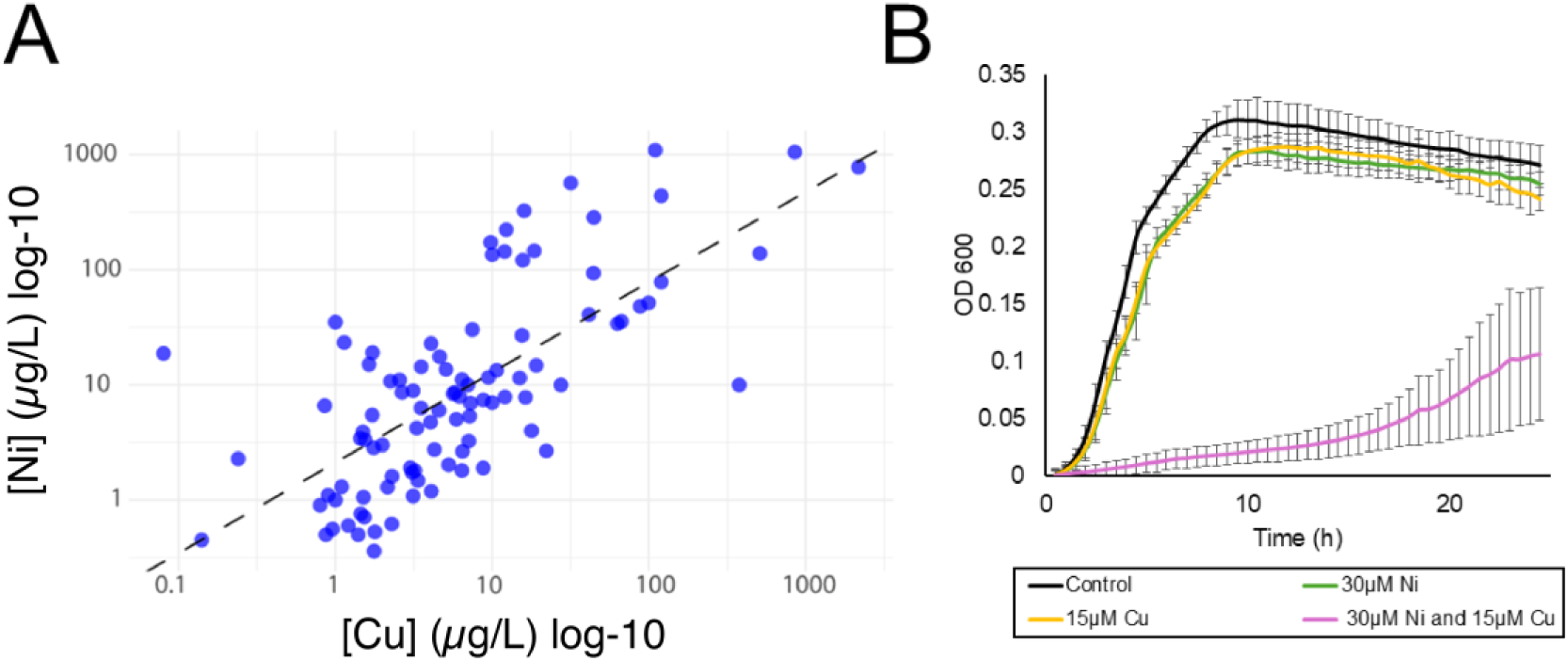
(A) Log–log correlation between Cu and Ni concentrations (µg/L) in global river and lake water bodies based on compiled environmental monitoring data from 1972 to 2017 (Zhou et al., 2020). The regression line for the equation [Ni] = 2.107[Cu]^0.784^ (R^2^ = 0.493, p = 1e-15) is shown as a dashed line. (B) Growth curves of *E. coli* under control conditions and treatment with 15 µM Cu, 30 µM Ni, and combined Ni and Cu exposure. Each point represents the average of 3 replicates and error bars represent ±SD.

### Synergistic toxicity of Ni and Cu

We next examined the growth of *E. coli* with a range of environmentally relevant concentrations of Ni^2+^ and Cu^2+^ to define our treatment conditions. Ni concentrations in freshwater have been reported between 0.006 and 157 µM (44–46) and Cu concentrations in freshwater range from 0.0126 to 432 µM (44–48). While Cu^2+^ is used for the amendments, it is reduced to Cu^1+^ intracellularly (49). In our experimental system, we will use “Ni” to refer to Ni^2+^ and “Cu” to refer to both the Cu^2+^ added to the cultures and the intracellular Cu^1+^, with distinction between the two made as needed. Our growth medium was modified from Rathnayake et al. and utilizes minimal nutrient concentrations, such as phosphate, to maximize free metal ion availability (35). As shown in **Supplementary Figure S1**, the growth of *E. coli* treated with 20 µM Ni and 10 µM Cu was comparable to the untreated control, indicating minimal toxicity. In contrast, treatment with 40 µM Ni and 20 µM Cu resulted in a significant reduction in bacterial growth. Based on these observations, an intermediate concentration of 30 µM Ni and 15 µM Cu was selected for further experiments. This combination was chosen to avoid lethality while still exerting a significant inhibitory effect on bacterial growth. We treated *E. coli* with either 30 µM Ni or 15 µM Cu, individually. These treatments resulted in minimal growth defects **(Fig. 1B)**. However, when the two metals were added in combination at the same concentrations (30 µM Ni and 15 µM Cu), a significant decrease in *E. coli* growth occurred **(Fig. 1B)**, suggesting that the toxicity of Ni and Cu in combination is synergistic, as described in older studies (50,51).

To characterize the cellular response to the combined Ni and Cu treatment, we conducted transcriptomic and metabolomic analyses of *E. coli* cultures treated with Ni (30 µM), Cu (15 µM), or their combination (30 µM Ni and 15 µM Cu). Untreated cultures served as controls. Changes in gene expression for each metal treatment were determined relative to the control after two hours of exposure during mid-log growth. In total, there were 512 differentially expressed genes across the three treatments **(Suppl. Table S2)**, of which 360 (70% of total) were uniquely differentially expressed in the combined metal treatment **(Fig. 2).** Untargeted metabolomic profiling revealed a similar trend **(Suppl. Fig. S2).** A total of 511 metabolites were differentially abundant across all treatments **(Suppl. Table S3).** Of these, 182 (35.6%) metabolites were uniquely differentially abundant only under the combined metal treatment. Interestingly, many of the differentially abundant metabolites (41.4%) were shared between the combined treatment and the Cu treatment alone. Overall, these findings highlight a synergistic interaction at both the phenotypic level **(Fig. 1B)** and the molecular level **(Fig. 2)**. We next analyzed the cellular systems that are differentially regulated during the combined Ni and Cu treatment to better understand [1] their mechanism of synergistic toxicity and [2] the cellular acclimation to this complex form of stress.

**Figure 2.**
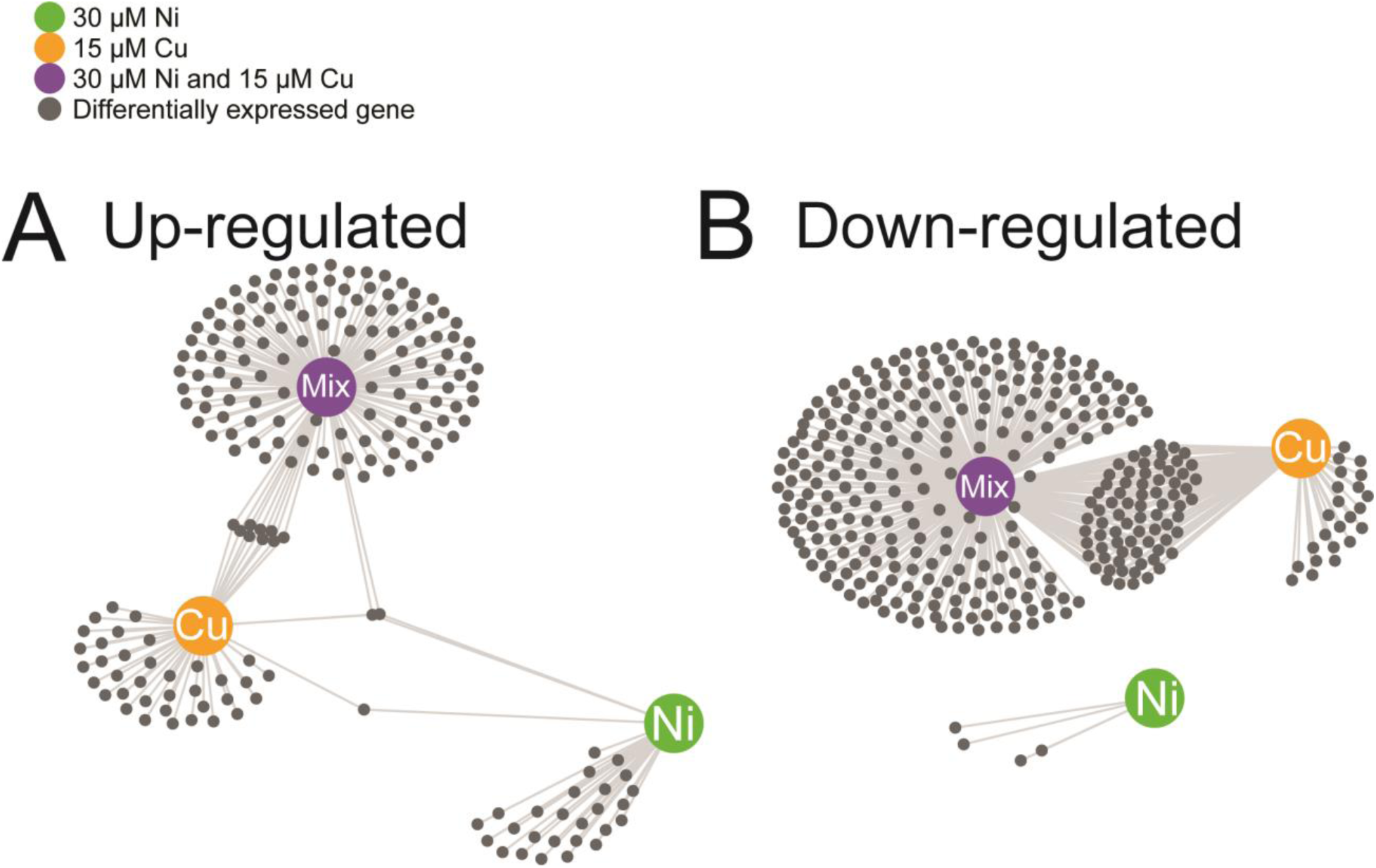
Transcriptomic analysis of *E. coli* reveals gene expression changes induced by 15 µm Cu, 30 µm Ni, and their combination. The network diagram shows differentially expressed (small, gray nodes) genes under each metal treatment (large, colored nodes). Edges connect treatment notes to gene nodes. (Panel A: upregulated genes; Panel B: downregulated genes). For each condition, three replicates were performed.

### ROS are not involved in the synergistic toxicity of Ni and Cu

Both Ni and Cu have been proposed to exert their cytotoxic effects, at least partially, through reactive oxygen species (ROS)-dependent mechanisms. Cu generates superoxide and hydroxyl radicals via Fenton chemistry *in vitro* (52). By itself, Ni is a poor free radical generator (53); however, when complexed by ligands such as cysteine (54), histidine (55), or oligopeptides (56), Ni also generates hydroxyl radicals *in vitro*. As our experiments were performed under aerobic growth conditions, we first considered whether ROS play a role in the synergistic toxicity of the two metals. However, we observed only a minimal transcriptional response from the “oxidative stress regulator” (OxyR) regulon under the three metal exposure conditions (57,58) **(Suppl. Table S4)**. We also monitored intracellular ROS levels in metal-treated cultures with the probe 2’,7’-dichlorodihydrofluorescein diacetate (H_2_DCFDA). In its intracellular, deacetylated form dichlorofluorescin (DCFH), the probe is responsive to hydrogen peroxide, hydroxyl radical, and peroxyl radical and is oxidized to the fluorophore dichlorofluorecsein (DCF) (59). The positive control (30% H₂O₂) caused an increase in ROS over time. However, treatment with either Ni, Cu, or their combination did not result in a significant increase in fluorescence compared to the untreated negative control **(Fig. 3).** From these data, we concluded that other molecular mechanisms are more important in the synergistic toxicity of Ni and Cu.

**Figure 3.**
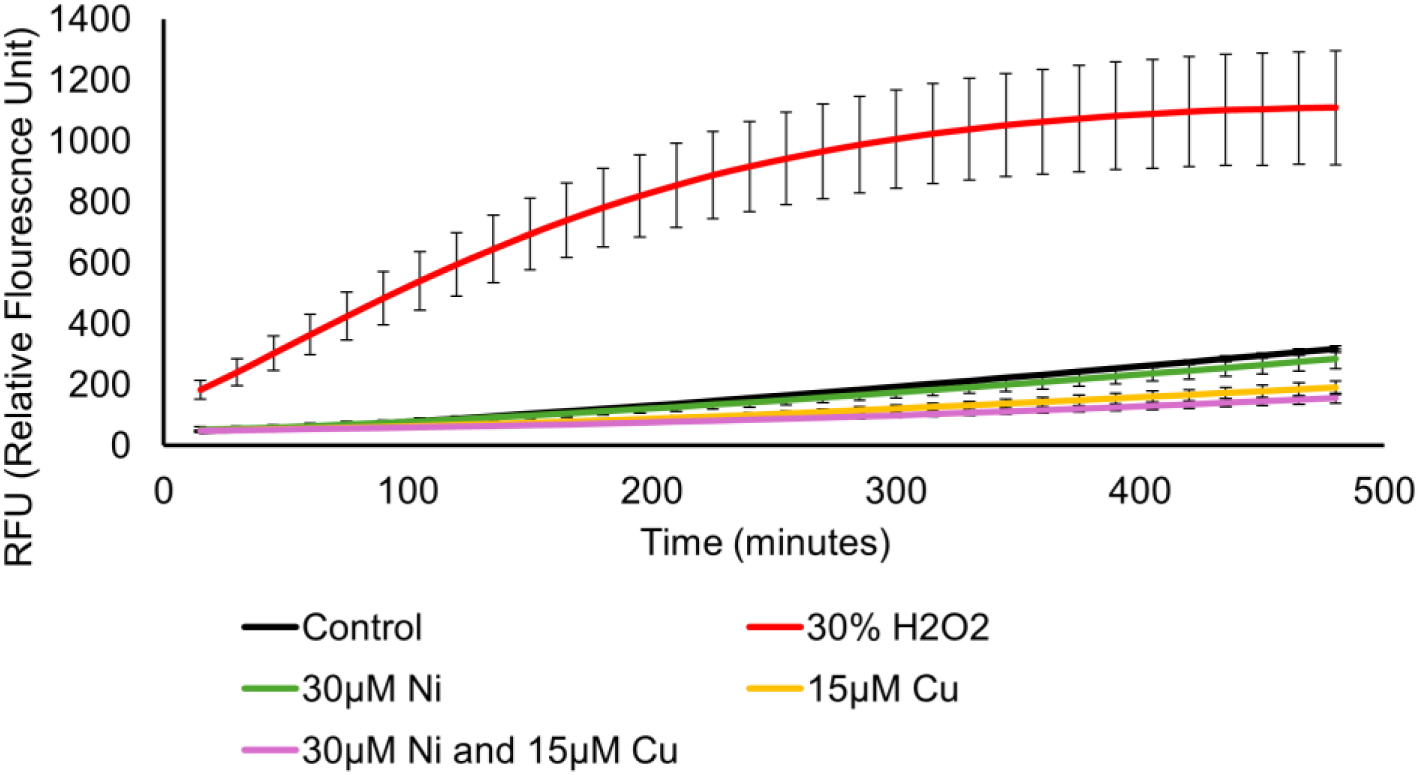
Measurement of intracellular ROS levels in *E. coli* following treatment with Ni, Cu, or combined metal stress. 30% hydrogen peroxide (H2O2) was used as a positive control. The “control” condition was untreated and serves as a negative control to establish the background level of fluorescence in the cultures. Each point represents the average of 3 replicates and error bars represent ±SD.

### Ni and Cu stress dysregulates histidine metabolism

We observed upregulation of the *his* operon (except *hisI*) during the combined Ni and Cu exposure, but not during individual metal treatments **(Fig. 4A)**. Metabolomics revealed elevated intracellular histidine concentrations during both the combined metal stress and Cu stress **(Fig. 4B)**. Addition of histidine to the growth medium rescued the growth defect from the combined metal exposure **(Fig. 4C)** and eliminated the minor growth inhibitionof the single metal exposures **(Suppl. Fig. S3).** Histidine forms coordinate complexes with Ni^2+^ (60) as well as Cu^1+^ and Cu^2+^ (61,62). With the exogenous histidine, extracellular metal chelation is likely occurring within the growth medium (*i.e.,* reducing the metal bioavailability (35,63)), but the same chemistry is expected to occur intracellularly.

**Figure 4.**
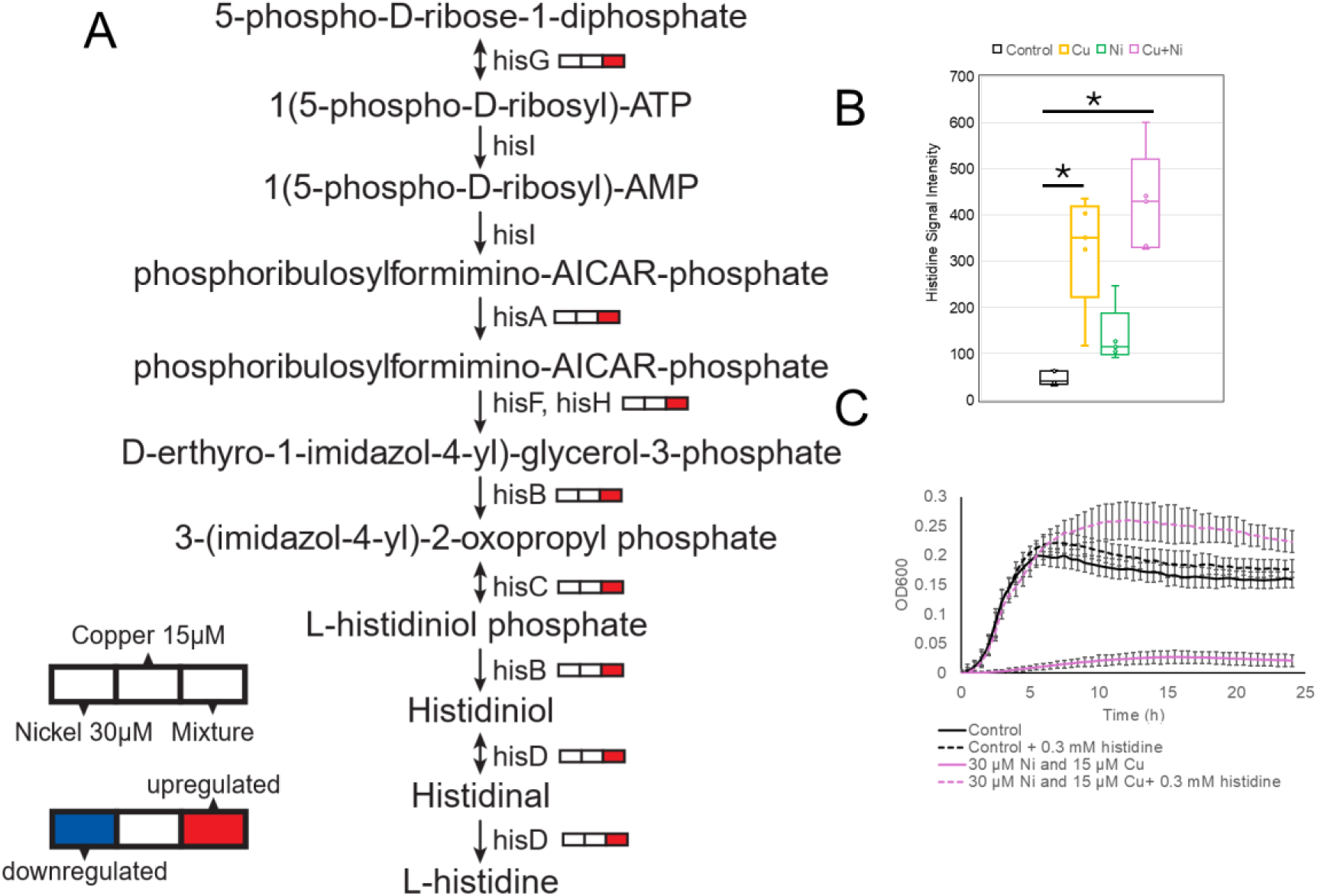
Histidine biosynthesis, metabolite levels, and functional rescue under metal stress. (A) Schematic of the histidine biosynthesis pathway in *E. coli*, highlighting genes that are differentially expressed following treatment with 30 µM Ni, 15 µM Cu, or a combination of both metals. (B) Intracellular histidine levels measured by metabolomics after exposure to individual metals or the combined treatment, relative to the untreated control. (C) Growth response of *E. coli* under combined metal stress (30 µM Ni + 15 µM Cu) with or without 0.3 mM histidine supplementation, compared to controls with and without histidine. Each point represents the average of 3 replicates and error bars represent ±SD.

Transcription of the *his* operon is regulated by an attenuation mechanism that is sensitive to His-tRNA^His^ levels, which reflect intracellular histidine concentrations (64). We propose that histidine complexation with Ni and Cu depletes free histidine, derepressing the operon and enhancing histidine biosynthesis (65). Since no additional Ni or Cu are added to the system, the newly synthesized histidine accumulates within the cytosol **(Fig. 4B).** In Cu-only conditions, histidine accumulation still occurred without operon induction, suggesting enzymatic feedback inhibition of HisG (ATP phosphoribosyltransferase) may be relieved by histidine binding to Cu (66). Complexation of intracellular histidine by Cu may be sufficient to relieve this feedback inhibition but not enough to relieve the transcriptional attenuation of the *his* operon. A similar, though nonsignificant, trend was observed for the Ni-exposed cultures **(Fig. 4B).** We propose that the combined effect of Ni and Cu on histidine biosynthesis could be understood as the cumulative result of the complexation of both metals with intracellular histidine.

Given that exogenous histidine protects against the combined Ni and Cu toxicity, it was attractive to speculate that increased expression of the *his* operon and accumulation of intracellular histidine is important in the cellular response to this combined stress (67,68). However, histidine biosynthesis mutant strains *ΔhisG* (ATP phosphoribosyltransferase) and *ΔhisD* (histidinol dehydrogenase) had similar growth phenotypes as the wild-type strain under all exposure conditions **(Suppl. Fig. S4)**, indicating that the observed biosynthetic changes are likely a chemical consequence of intracellular metal complexation, not an acclimatization response. This is in line with prior studies that also reported increased expression of the *his* operon in other bacteria in response to high concentrations of either Ni or Cu (69–71).

### Modulation of sulfur metabolism in the response to combined Ni and Cu stress

Upon exposure to the combined Ni and Cu treatment, we observed significant upregulation of multiple genes involved in sulfate assimilation **(Fig. 5A),** including *cysN* (sulfate adenylyltransferase subunit 1), *cysC* (adenylyl-sulfate kinase), *cysD* (sulfate adenylyltransferase subunit 2), *cysH* (phosphoadenosine phosphosulfate reductase), *cysI* (sulfite reductase [NADPH] hemoprotein β-subunit)*, cysJ* (sulfite reductase [NADPH] flavoprotein α-subunit), and *cysK* (cysteine synthase A). This response did not occur with either the Ni or Cu treatments. Similarly, we observed the upregulation of *cysA* (ATP-binding component of the CysPUWA sulfate/thiosulfate ABC transporter) and *sbp* (sulfate/thiosulfate ABC transporter periplasmic binding protein) under the combined Ni and Cu treatment only **(Suppl. Table S2)** (72). This pathway is essential for assimilating inorganic sulfur into cysteine, a sulfur-containing amino acid (73). Consistent with our transcriptional data, supplementing the culture medium with 20 mM sulfate partially rescued growth during the combined metal stress **(Fig. 5D).** In contrast, the addition of 20 mM sulfate to cultures treated with either Cu or Ni individually had no significant effect on growth **(Suppl. Fig. S5).**

**Figure 5:**
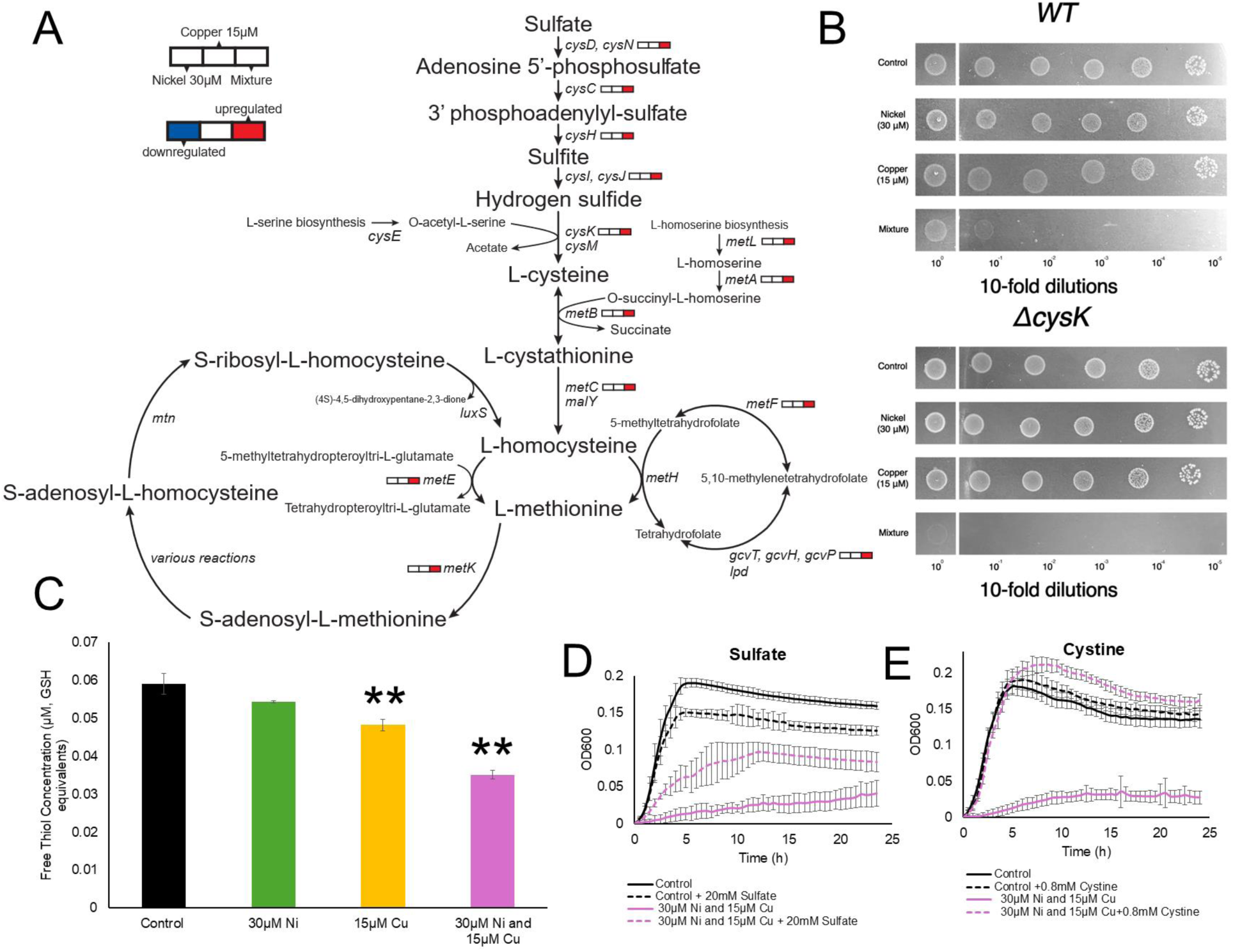
Sulfur assimilation, cysteine, and methionine metabolism are impacted by metal stress in *E. coli.* (A) Transcriptomic modulation of the sulfur assimilation and methionine biosynthesis pathways in *E. coli* during exposure to 30 µM Ni, 15 µM Cu, and their combination (Ni+Cu), relative to untreated controls. (B) Spot dilution assay comparing growth of *ΔcysK* mutant and wild-type *E. coli* under control conditions and metal treatments. Experiments were performed a minimum of two times, with one representative trial shown. (C) Quantification of total intracellular thiol levels in cells exposed to Ni, Cu, and Ni+Cu treatments. ** (p < 0.01); *** (p < 0.001). (D) Impacts of sulfate (20 mM) supplementation on growth under combined metal stress (30 µM Ni + 15 µM Cu). (E) Effect of 0.8 mM cystine supplementation on growth under combined metal stress. For panels C-E, each point represents the average of 3 replicates and error bars represent ±SD.

We hypothesized that this sulfur demand might stem from an increased need for cysteine and/or cysteine-derived compounds like glutathione or methionine during the combined metal stress. Indeed, we observed a decrease in the total intracellular thiol concentrations during the exposure to the combined metal stress **(Fig. 5C).** Although glutathione supplementation improved growth under the combined metal stress (and Cu stress) **(Suppl. Fig. S6)**, *gshA* (glutamate-cysteine ligase) and *gshB* (glutathione synthetase) (74) expression remained unchanged across all conditions **(Suppl. Table S2)**, and their deletion improved growth under the combined metal stress **(Suppl. Fig. S7)**. We speculate that glutathione amendments likely rescued growth, in part, through the extracellular chelation of the Cu^2+^ (75,76).

Methionine biosynthesis genes were upregulated **(Fig. 5A)** and intracellular methionine levels increased under the combined metal stress while levels of the pathway intermediate cystathionine decreased **(Suppl. Fig. S8A)**. Methionine supplementation (7 mM) also significantly rescued growth under the combined metal stress but did not affect growth under the single metal stressors **(Suppl. Fig. S8B)**. However, like glutathione, the methionine biosynthesis mutant strains *ΔmetA, ΔmetB,* and *ΔmetE* outperformed wild type strain **(Suppl. Fig. S9),** suggesting that methionine overproduction may be maladaptive during growth under the combined metal stress.

We next tested whether the sulfur limitation stress response was due to increased cysteine demand. We supplemented the metal-stressed cultures with 0.8 mM cystine, the disulfide form of cysteine that is reduced to cysteine intracellularly (77). Cystine completely alleviated the growth inhibition observed in the cultures treated with Ni and Cu, as well as Cu or Ni individually **(Fig. 5E) (Suppl. Fig. S10).** In our transcriptome data, we noted the up-regulation of genes encoding two cystine transporters during the combined metal stress: TcyP and the TcyJ subunit of the TcyJLN transporter **(Suppl. Table S2).** Considering the increased expression of the cysteine synthase A gene (*cysK*) during the combined metal exposure **(Fig. 5A),** we next tested the growth of a *ΔcysK* mutant under our different exposure conditions. *ΔcysK* had comparable growth to the wild-type strain under the individual Cu or Ni exposures **(Fig. 5B).** In contrast, *ΔcysK* had weaker growth with Cu and Ni compared to the wild-type strain **(Fig. 5B**). Thus, cysteine biosynthesis is critical for survival under the combined metal stress. Additionally, since our growth medium supplies amino acids in the form of yeast extract, we propose that the disruption of the methionine or glutathione biosynthesis pathways improved growth **(Suppl. Figs. S7 and S9)** during the combined metal stress by rerouting cysteine to an unknown, growth-promoting pathway(s) active under these conditions.

### Iron-sulfur clusters are targeted by the combined metal stress

One plausible sink for this cysteine is [Fe-S] cluster assembly since cysteine is the sulfur donor for [Fe-S] cluster biogenesis (2). Consistent with this model, we observed increased expression of genes involved in iron-sulfur [Fe-S] cluster assembly under combined metal stress **(Fig. 6A).** *E. coli* utilizes two main systems for Fe-S cluster biosynthesis: [1] the ISC (Iron-Sulfur Cluster) system, which functions as the primary housekeeping pathway and [2] the SUF (Sulfur Utilization Factor) system, which is induced under stress conditions (78). Genes associated with the ISC system, including, *iscS* (cysteine desulfurase), *iscU* (scaffold protein) and *iscA* (iron-sulfur cluster carrier protein), were upregulated in response to the metal mixture, but not under individual Ni or Cu treatments. In contrast, the SUF system genes were not differentially expressed under any condition. We also observed up-regulation of *erpA* (Essential Respiratory Protein A) under the combined metal stress and Cu treatment. ErpA plays a crucial downstream role in delivering preassembled Fe-S clusters to apoprotein targets, particularly those involved in essential cellular processes like respiration and central metabolism (79).

**Figure 6:**
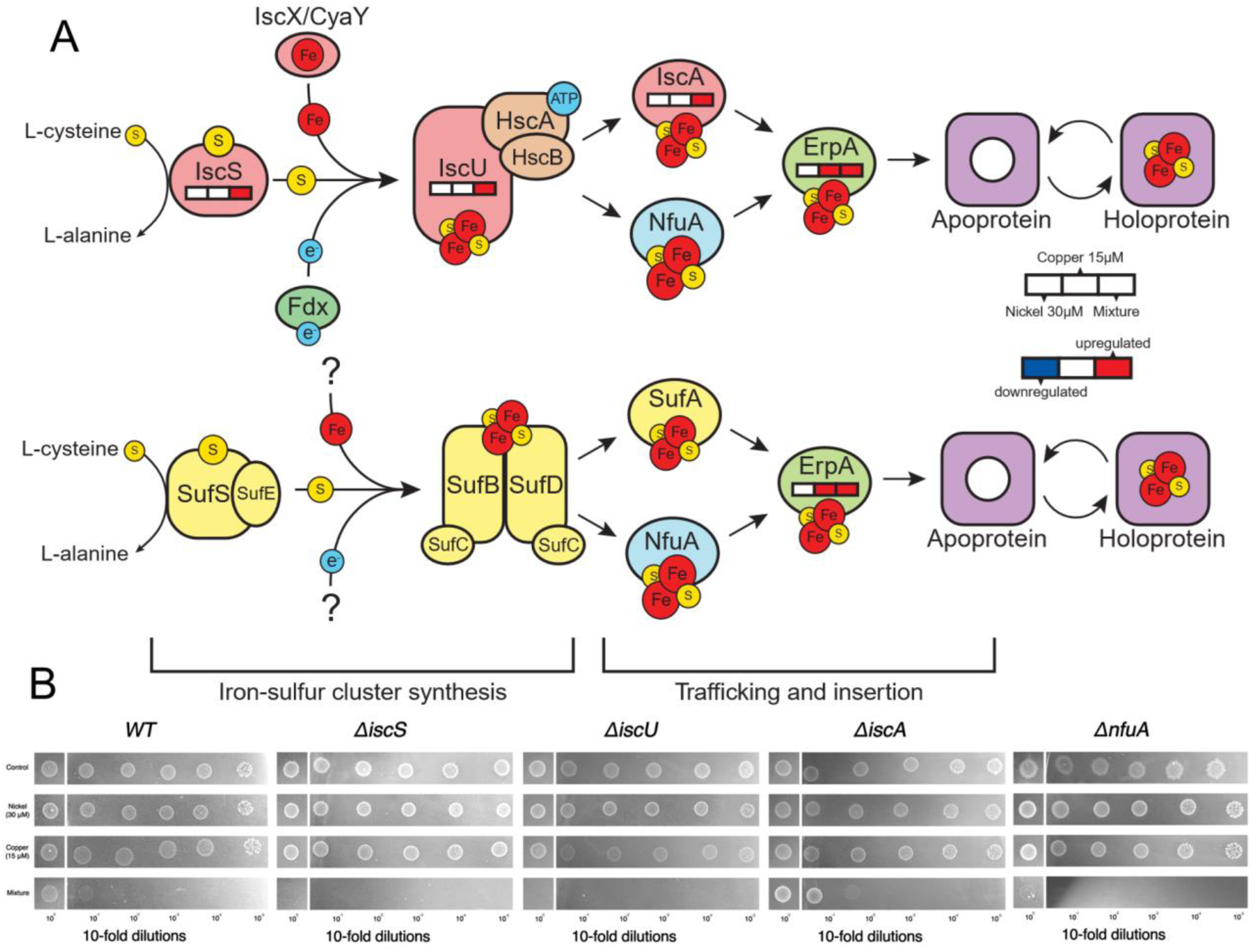
Iron–sulfur cluster biogenesis pathway and mutant phenotypes under metal stress conditions. (A) Schematic representation of the iron–sulfur (Fe–S) cluster synthesis pathway in *E. coli*, including its trafficking and insertion into target proteins. Genes involved in this pathway are overlaid with transcriptomic data showing differential expression in response to 30 µM Ni, 15 µM Cu, and combined Ni+Cu treatment. (B) Spot dilution assay showing the growth of wild-type *E. coli* and deletion mutants *(ΔiscS, ΔiscU, ΔiscA*, and *ΔnfuA*) under control and metal stress conditions. Serial 10-fold dilutions (10⁻^0^ to 10⁻⁵) were spotted on MES agar plates containing no metal (control), 30 µM Ni, 15 µM Cu, or both metals combined. Experiments were performed a minimum of two times, with one representative trial shown.

We next tested the growth of Fe-S cluster assembly and trafficking mutants with the different metal exposures. *ΔiscS* and *ΔiscU* strains were unable to grow under combined metal exposure, indicating that the core assembly machinery of the ISC system is essential under this condition **(Fig. 6B).** In contrast, the *ΔiscA* mutant exhibited growth comparable to the wild type **(Fig. 6B)**, consistent with the known redundancy of auxiliary ISC components (80–82). *ΔnfuA* had weaker growth with the metal combination relative to the wild-type strain **(Fig. 6B)**. This aligns with recent findings that NfuA interacts with ErpA to transfer Fe-S clusters from the ISC and SUF scaffolds to essential respiratory and other metabolic proteins (83,84). *erpA* itself is essential in *E. coli* and cannot be deleted (83). We also confirmed that the SUF system was not required for growth under the combined metal stress. Under the combined metal stress, both *ΔsufD* and *ΔsufS* had, surprisingly, enhanced growth while *ΔsufA* had similar growth relative to the wild-type strain **(Suppl. Fig. S11)**. None of the mutant strains had altered growth under any of the other conditions tested **(Fig. 6B, Suppl. Fig. S11**).

The observation that SUF was not essential for growth under the combined metal stress was surprising as the SUF system is typically induced under stress conditions where the ISC system is impaired, specifically oxidative damage or iron (Fe) limitation (85). However, this finding was consistent with our earlier observation that intracellular ROS levels did not increase significantly under any metal exposure **(Fig. 3).** Fe^2+^ supplementation also failed to rescue growth **(Suppl. Fig. S12)**, nor did we observe significant de-repression of the Fur regulon (86) under the combined metal exposure. In fact, expression of the *feoABC* operon, encoding the Fe^2+^ transporter, was decreased during the combined metal exposure (87) **(Suppl. Table S2)**. Thus, sulfur—rather than Fe—appears to be the limiting nutrient for Fe-S biosynthesis under combined metal stress. This contrasts with our prior findings in *B. cereus*, where a more complex metal mixture led to a broad Fe starvation response (30).

The *iscRSUA-hscBA-fdx (isc)* operon and *erpA* are both regulated by the transcription factor IscR. When bound to a [2Fe-2S] cluster, IscR represses the expression of these genes (88). Prior studies in *E. coli* suggest that loss of the bound [2Fe-2S] leads to de-repression of these genes and occurs through two mechanisms [1] decreased synthesis of Fe-S clusters (89) and [2] damage to extant Fe-S clusters (90). We propose a model where both might occur during the combined metal stress, leading to increased expression of the ISC system as well as *erpA* **(Fig. 6A).** The divalent metal cations cobalt (Co^2+^), zinc (Zn^2+^), and cadmium (Cd^2+^) bind to the active site of the *E. coli* IscU scaffold protein (91), whereas Cu^1+^ does not bind to IscU (92). This selectivity is governed by the coordination geometry preferences of the IscU active site (91). It is conceivable that Ni^2+^, with its similar coordination chemistry to Co^2+^ (93), selectively binds to IscU over the Cu^1+^. Furthermore, due to its placement within the Irving-Williams series, Ni^2+^ would also out-compete cellular Fe^2+^ for IscU binding (94). Binding of IscU to transition metals other than Fe^2+^ deactivates the enzyme, inhibiting *de novo* Fe-S cluster biosynthesis (95,96). While Cu^1+^ does not bind IscU, Cu^1+^ does damage existing Fe–S clusters by directly displacing Fe through interaction with coordinating cysteine thiolates, deactivating Fe-S cluster-containing enzymes (97). In contrast, *in vitro* experiments have suggested that Ni^2+^ does directly not alter the activity of Fe-S cluster-containing enzymes (98).

According to our model, during Cu exposure, Fe-S cluster degradation can be countered by maturation of new Fe-S by a functional ISC system. We see some evidence for this in our transcriptome data with increased expression *erpA* during Cu exposure **(Fig. 6A).** During Ni exposure, maturation of new Fe-S clusters is less important if there is minimal damage to existing clusters—supported by the lack of transcriptional changes related to Fe-S cluster biosynthesis under these conditions. We propose that the two metals synergize in their toxicity by [1] damaging the existing Fe-S clusters within enzymes and then [2] inhibiting biosynthesis of new clusters. This model provides further context to our observation that sulfur is the limiting nutrient for Fe-S biosynthesis under combined metal stress, rather than Fe. Metal-induced damage to Fe-S clusters increases intracellular Fe^2+^ concentrations (98–100), which would explain the decreased expression of the Fur-regulated *feoABC* during both the combined metal stress and the individual Cu stress **(Suppl. Table S2).** Dysregulation of Fe-S cluster metabolism by the combined metal stress also puts a significant strain on cellular sulfur metabolism in two ways. First, as we noted earlier, cysteine is the sulfur donor for ISC (78). Second, assimilatory sulfate reduction to cysteine is dependent upon Fe-S cluster metabolism as the sulfite reductase (CysJI) has multiple Fe-S cluster cofactors. Damage to Fe-S clusters is known to reduce sulfite reductase activity (101). This would explain our observation that high concentrations of sulfate (20 mM) only partially rescue growth during the combined metal stress while cystine at a lower concentration (0.8 mM cystine, or 1.6 mM cysteine) fully rescues growth under the same condition **(Fig. 5D, E).** Decreased metabolic flux through the sulfur assimilation pathway may also have caused the increase in expression of the methionine biosynthesis pathway under the combined metal stress **(Fig. 5A).** In *Salmonella enterica* it was reported that Co^2+^ damages Fe-S clusters, leading to decreased sulfite reductase activity, which resulted in a methionine auxotrophic requirement (101,102)

### Reprogramming of the TCA Cycle

Fe-S clusters have essential roles in electron transfer, central metabolism, and gene regulation, making them critical to cellular functioning (2). Therefore, we considered the downstream consequences of Fe-S cluster disruption under combined metal stress. Transcriptomic analysis revealed modulation of the tricarboxylic acid (TCA) cycle and related pathways only under the combined metal treatment **(Fig. 7).** Several enzymes of the TCA cycle (*i.e.,* aconitase (103), succinate dehydrogenase (104), fumarase (105)), are dependent on Fe-S clusters for their activities. For example, *fumC* (fumarase C), encoding an alternative, [Fe-S]-independent fumarase (105), was upregulated only in the combined Ni and Cu condition **(Fig. 7)**. The small RNA *spf* (small regulatory RNA Spot 42), which inhibits the translation of key TCA cycle genes— *gltA* (citrate synthase) and *sdhCDAB* (succinate dehydrogenase) **(Fig. 7)** (106–108)— was also downregulated under both Cu and combined metals treatments. Decreased expression of *spf* would relieve translational repression of key TCA cycle enzymes, promoting sustained flux through the TCA cycle to support energetic and biosynthetic demands under combined metal stress.

**Figure 7:**
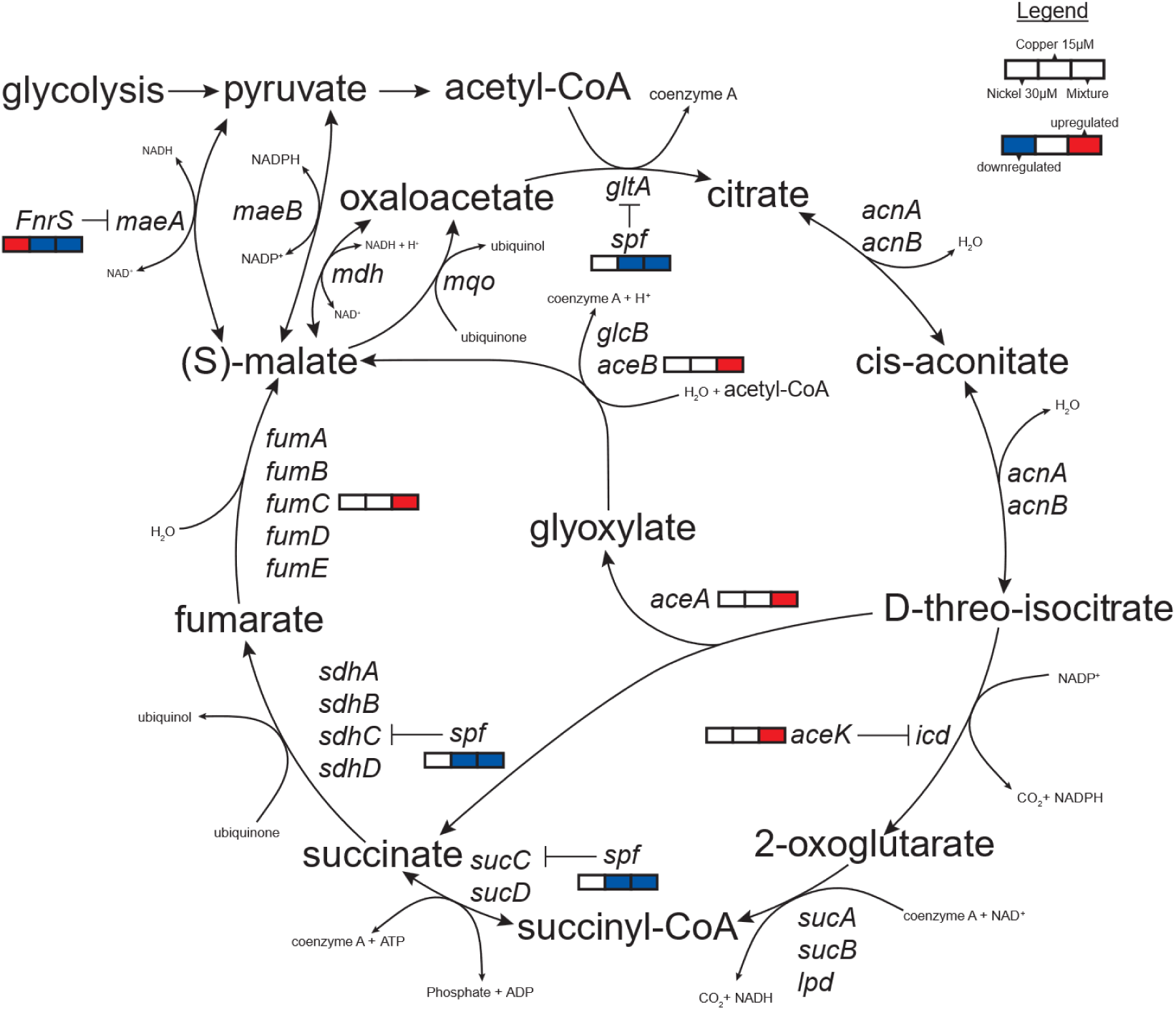
Changes in the expression of genes encoding TCA cycle and glyoxylate shunt enzymes and small RNAs (*fnrS* and *spf*) during the metal exposures. Genes involved in this pathway are overlaid with transcriptomic data showing differential expression in response to 30 µM Ni, 15 µM Cu, and the combined Ni+Cu treatment.

Genes of the glyoxylate shunt—*aceA* (isocitrate lyase), *aceB* (malate synthase A), and *aceK* (isocitrate dehydrogenase kinase / isocitrate dehydrogenase phosphatase)—were significantly upregulated only under the combined metal stress**(Fig. 7)**. This shunt bypasses the CO₂-releasing steps of the TCA cycle, allowing *E. coli* to conserve carbon and redirect intermediates toward anabolic pathways (109–111). AceA performs the first step of the shunt, the cleavage of isocitrate to succinate and glyoxylate, while AceB converts glyoxylate to malate. AceK is the regulatory switch that turns on or off the activity of isocitrate dehydrogenase, controlling the partitioning of carbon between the TCA cycle and glyoxylate shunt. While glyoxylate concentrations were increased during the combined metal exposure, it did not reach the level of significance **(Suppl. Fig. S13)**. Malate appears to be maintained at a low level in the cells as it was not detected under any condition. Succinic acid was significantly increased during the combined metal exposure **(Suppl. Fig. S13)**. This could be attributed to the increased flux through the glyoxylate shunt, decreased activity of the Fe-S-cluster enzyme succinate dehydrogenase (112), or a combination of the two.

The glyoxylate shunt has been implicated in the response to stressors including hypoxia (113), ROS (111,114), antibiotics (115,116), and Fe limitation (117,118). During Fe limitation, shifting carbon flux through the glyoxylate shunt allows cells to bypass the steps of the TCA cycle reliant upon the heavily Fe-cofactor-dependent electron transport chain (ETC). The same logic could be applied to multi-metal stress as Complex I of the *E. coli* aerobic ETC, required for NADH re-oxidation, has multiple Fe-S cluster cofactors. However, it is also possible that conservation of carbon via the glyoxylate shunt is required to support the *de novo* synthesis of amino acids like histidine, cysteine, and methionine, whose biosynthetic pathways are up-regulated during the combined metal exposure **(Figs. 4, 5A).** We next tested whether this shift from the full TCA cycle to the glyoxylate is adaptive by examining the growth of the mutant strains *ΔaceB, ΔaceA,* and *ΔaceK* under the different metal exposures **(Suppl. Fig. S14)**. *ΔaceB* and *ΔaceA* had similar phenotypes of the wild-type strain while *ΔaceK* had a more severe growth defect under the combined metal stress. Under all other conditions, growth the three mutant strains grew similarly to the wild-type strain. These data suggest that the regulatory switch between the full TCA cycle and the glyoxylate shunt by AceK is critical in the cellular response to the metal mixture stress.

The post-transcriptional regulatory small RNA *fnrS* showed condition-specific expression: it was downregulated under Cu stress and the combined metal exposures but upregulated during Ni exposure alone. One *fnrS* target, *maeA* (malate dehydrogenase), links the TCA cycle with pyruvate metabolism **(Fig. 7)** (119,120). Reduced *fnrS* under combined stress may increase maeA translation, boosting malate-to-pyruvate flux and enhancing carbon routing flexibility. *fnrS* is up-regulated by Fnr (121), a global regulator of the aerobic-anaerobic transition that is itself activated by a [4Fe-4S] cluster (122,123). This cluster is sensitive to damage from oxygen (124) or nitric oxide (125) and maybe also from Cu during the combined metal stress. While our experiments were performed under aerobic growth conditions, during mid-to-late exponential growth—the period captured by our transcriptome experiments—oxygen typically becomes depleted in shaken flask cultures of *E. coli*, leading to increased expression of the Fnr regulon (90,126). This exponential, aerobic-to-anaerobic transition is apparently lost during the combined metal stress. Consistent with Fnr inactivation, multiple Fnr-regulated genes showed reduced expression under combined metal stress, including *dmsA* (DMSO reductase subunit A)*, nikC* (nickel ABC transporter membrane subunit)*, yfiD* (stress-induced alternate pyruvate formate-lyase)*, narGHIJ* (respiratory nitrate reductase)*, ydfZ* (selenoprotein YdfZ)*, nirBDC* (nitrite reductase)*, nrfA* (assimilatory nitrate reductase)*, caiF* (transcriptional activator CaiF)*, narK* (nitrate:nitrite antiporter), *frdABC* (fumarate reductase)*, hcp* (nitric oxide reductase)*, dcuB* (anaerobic C4 dicarboxylate transporter)*, dcuC* (anaerobic C4 dicarboxylate transporter), *ynfK* (dethiobiotin synthetase), and *yjiM* (dehydratase) (127). Many of these same genes were also downregulated during Cu exposure (**Suppl. Table S2).** These expression patterns support a model where, during combined metal stress, the Cu damages the Fnr cluster, repressing the Fnr regulon.

Altogether, these transcriptional changes suggest that co-exposure to Ni and Cu induces a broad metabolic reprogramming involving both enhancement of key TCA cycle functions, activation of the glyoxylate shunt, and inhibition of pathways involved in adaptation to anaerobiosis. We speculate that many of these observed changes likely link back to dysregulation of Fe-S cluster metabolism; however, future work is required to confirm this model.

## CONCLUSIONS AND ENVIRONMENTAL SIGNIFICANCE

Freshwater concentrations of Cu are typically below (44,46) regulatory thresholds set by the World Health Organization (31 µM) and the US Environmental Protection Agency (USEPA 20 µM) (128). For example, a recent survey of 179 freshwater bodies found that only three sites exceeded this USEPA limit (44), with a median Cu concentration of 0.11 µM. Even in contaminated environments, Cu concentrations often do not surpass action levels. For instance, in an aquifer impacted by improper heavy metal waste disposal, Cu concentrations were reported at 0.005 to 15 µM (46). In a stream near a Cu mine, Cu concentrations were no higher than 16 µM (129) However, Cu rarely exists in isolation of other metals (44). It frequently co-occurs with other metals such as Ni **(Fig. 1A).** We propose that Ni, which is positively correlated with Cu across environmental datasets **(Fig. 1A),** amplifies Cu toxicity in bacteria by potentiating Cu-induced disruption of Fe-S cluster metabolism. Fe-S clusters are central to all forms of microbial metabolism including generation of biosynthetic precursors (104,130), amino acid biosynthesis (97,131), aerobic respiration (132), nitrate respiration (133), sulfate respiration (134), fermentation (135), nitrogen-fixation (136), methanogenesis (137), acetogenesis, and carbon fixation. While our study uses *E. coli* as a model, this vulnerability likely extends across diverse prokaryotic and eukaryotic microbes. Understanding the synergistic toxicity of co-occurring metals like Ni and Cu is critical for accurately predicting microbial responses to metal pollution, a growing threat to ecosystem function in the Anthropocene (138).

## DATA AVAILABILITY

Raw reads for the transcriptomic data are deposited in the Sequence Read Archive under accession numbers SRR34735813-SRR34735824.

## ACKNOWLEDGEMENT

This work was supported by a Faculty Fellows Program award to JLG from the Syracuse Center of Excellence in Environmental and Energy Systems, which is funded by the Empire State Development’s Division of Science, Technology, and Innovation (NYSTAR). Additional funding was provided by the award from the SUNY Center for Applied Microbiology to JLG and start-up funds for JLG from SUNY ESF. RAI received financial support from the SUNY Chancellor’s Summer Research Excellence Fund. We thank Felicity Goetz for their assistance in documenting the mutant growth in the spot-plating assays.

